# Duplications drive diversity in *Bordetella pertussis* on an underestimated scale

**DOI:** 10.1101/2020.02.06.937284

**Authors:** Jonathan S. Abrahams, Michael R. Weigand, Natalie Ring, Iain MacArthur, Scott Peng, Margaret M. Williams, Barrett Bready, Anthony P. Catalano, Jennifer R. Davis, Michael D. Kaiser, John S. Oliver, Jay M. Sage, Stefan Bagby, M. Lucia Tondella, Andrew R. Gorringe, Andrew Preston

## Abstract

Bacterial genetic diversity is often described using solely base pair changes despite a wide variety of other mutation types likely being major contributors. Tandem duplications of genomic loci are thought to be widespread among bacteria but due to their often intractable size and instability, comprehensive studies of the range and genome dynamics of these mutations are rare. We define a methodology to investigate duplications in bacterial genomes based on read depth of genome sequence data as a proxy for copy number. We demonstrate the approach with *Bordetella pertussis*, whose insertion sequence element-rich genome provides extensive scope for duplications to occur. Analysis of genome sequence data for 2430 *B. pertussis* isolates identified 272 putative duplications, of which 94% were located at 11 hotspot loci. We demonstrate limited phylogenetic connection for the occurrence of duplications, suggesting unstable and sporadic characteristics. Genome instability was further described in-vitro using long read sequencing via the Nanopore platform. Clonally derived laboratory cultures produced heterogenous populations containing multiple structural variants. Short read data was used to predict 272 duplications, whilst long reads generated on the Nanopore platform enabled the in-depth study of the genome dynamics of tandem duplications in *B. pertussis*. Our work reveals the unrecognised and dynamic genetic diversity of *B. pertussis* and, as the complexity of the *B. pertussis* genome is not unique, highlights the need for a holistic and fundamental understanding of bacterial genetics.

## Introduction

*Bordetella pertussis* is a Gram-negative bacterium which is the main causative agent of the human respiratory disease whooping cough. *B. pertussis* has speciated from a *B. bronchiseptica*-like ancestor to become a host restricted pathogen (Diavatopoulos et al. 2005; Parkhill et al. 2003). This process has occurred primarily via genome reduction: the *B. bronchiseptica* genome is around 5.4Mbp whereas the *B. pertussis* genome is around 4.1Mbp, involving loss of over 1000 genes during speciation, and has been driven primarily by deletions arising from recombination between insertion Sequence (IS) elements (Preston et al. 2004; Parkhill et al. 2003). Genomes of *B. pertussis* strains include over 240 copies of IS*481*, with far fewer copies of IS*1663* and IS*1002*. Gene erosion in *B. pertussis* appears to be on-going and sporadic IS-mediated deletions and disruptions provide subtle differences in gene content between strains (King et al. 2010; Heikkinen et al. 2007; Caro et al. 2008), but there is little understanding of the effects.

Using the most popular metric of genetic diversity, single nucleotide polymorphisms (SNPs), *B. pertussis* is a species with extraordinarily low diversity leading to its description as a monomorph (Mooi 2010; Weigand et al. 2017). More detailed analyses of *B. pertussis* genome sequences have been limited by the inability to generate closed genome assemblies from short-read sequencing data, as the reads do not span IS*481* (1043 bp), and the assembly produces many contigs-consistently in excess of the number of IS*481* copies. Recent advances in long-read sequencing, notably by Pacific Biosciences (PacBio) and Oxford Nanopore Technologies (Nanopore), has enabled routine generation of closed genome assemblies for *B. pertussis* (Weigand et al. 2018a, 2016; Ring et al. 2018; Weigand et al. 2017; Bowden et al. 2016). Subsequent comparative analyses have revealed that intragenomic recombination between IS*481* causes genomic rearrangement and that a large number of different genome orders exist among circulating *B. pertussis* isolates (Weigand et al. 2017). The effect of rearrangement on *B. pertussis* phenotype remains unknown but moving genes between leading and lagging strands and to different locations in the chromosome would be expected to alter their expression (Price et al. 2005; Rocha and Danchin 2003). Likewise, transcription from IS element promoters can affect neighbouring genes, and different copies of IS*481* exhibit different transcriptional activities (Amman et al. 2018). Rearrangements that shuffle IS element-neighbouring gene combinations might, therefore, elicit changes in gene expression profiles both locally and genome-wide.

In addition to deletion and rearrangement, IS-mediated recombination can result in duplication. Twelve copy number variants (CNVs) in *B. pertussis* have been described and studies with sufficient genomic data have resolved them as tandem repeats (Caro et al. 2006; Dalet et al. 2004; Dienstbier et al. 2018; Heikkinen et al. 2007; Weigand et al. 2016, 2018a). Duplication of a region containing *cya*A (encoding adenylate cyclase-haemolysin) increased haemolytic activity and it was noted that this duplication was highly unstable (Dalet et al. 2004). These serendipitous observations suggest that CNVs are a poorly characterised contributor to genetic diversity among *B. pertussis*. However, to date there has been no systematic analysis of CNVs in *B. pertussis* and indeed systematic analysis of structural variants at the species level is rare for bacteria, although it is relatively common in eukaryotic organisms. In this study we sought to catalogue CNVs in *B. pertussis*, utilising publicly available genomic data, which is overwhelmingly derived from short-read sequencing platforms.

Among genomic data from 2430 *B. pertussis* isolates we found 191 which contained evidence of CNVs and identified that 94% of CNVs occur at 11 ‘hotspot’ loci. Some CNVs were very large, exceeding 300 kb in length. We reveal that some regions are present in multi-copy, and thus use the term copy number variant (CNV) rather than duplication. We contextualise this information using phylogenetics and find that strains containing similar CNVs are often distantly related, suggesting that CNVs at hotspot loci arise independently. Also, we confirm that laboratory grown populations of cells contain a mixture of copy numbers suggesting that CNV formation is a dynamic process, at least at some loci. Our study revealed novel genetic variation among *B. pertussis* isolates and provides a blueprint for investigation of CNVs in other bacteria, particularly those with high numbers of repeats.

## Results

The US Centers for Disease Control and Prevention (CDC) conducts routine and enhanced surveillance of pertussis, which includes whole genome sequencing of *B. pertussis* clinical isolates using the PacBio and Illumina platforms. Some of these data revealed increased read depth coverage localized to discrete genomic regions in some strains. Sequence data alone was incapable of resolving assembly of these regions but enzyme mapping of high-molecular weight DNA confirmed that the high read depth resulted from tandem CNVs. In total, genomes from 28 strains, including two used for the production of vaccines against pertussis, were identified which contained CNVs (Supplemental_Table_1) (Weigand et al. 2016). Some of these CNVs are large (>300kb), involving hundreds of genes. The accurate assembly of these genomes required manual resolution, using data from short read, long read and enzyme mapping sources. Using the manually resolved dataset as a benchmark, we sought to develop a prediction and screening tool to identify CNVs within the public repository of *B. pertussis* genome sequence data on the Sequence Read Archive (SRA) using a scalable and automated approach.

### Read depth as a proxy for copy number

We mapped short-read data from each query strain to a reference genome and used read depth as a proxy for copy number of genomic regions (Figure 1). If a strain contained two copies of a locus present at single copy in the reference genome, twice as many reads should be detected that map to that locus. Conversely, a gene deletion present in a query strain produces zero read depth at that locus in the reference. Since coverage depth fluctuates during whole genome sequencing due to a combination of biases and stochasticity (Ekblom et al. 2014; Loman et al. 2012), read depth coverage data was normalised and statistically analysed using the tool CNVnator (Abyzov et al. 2011).

**Figure 1.**
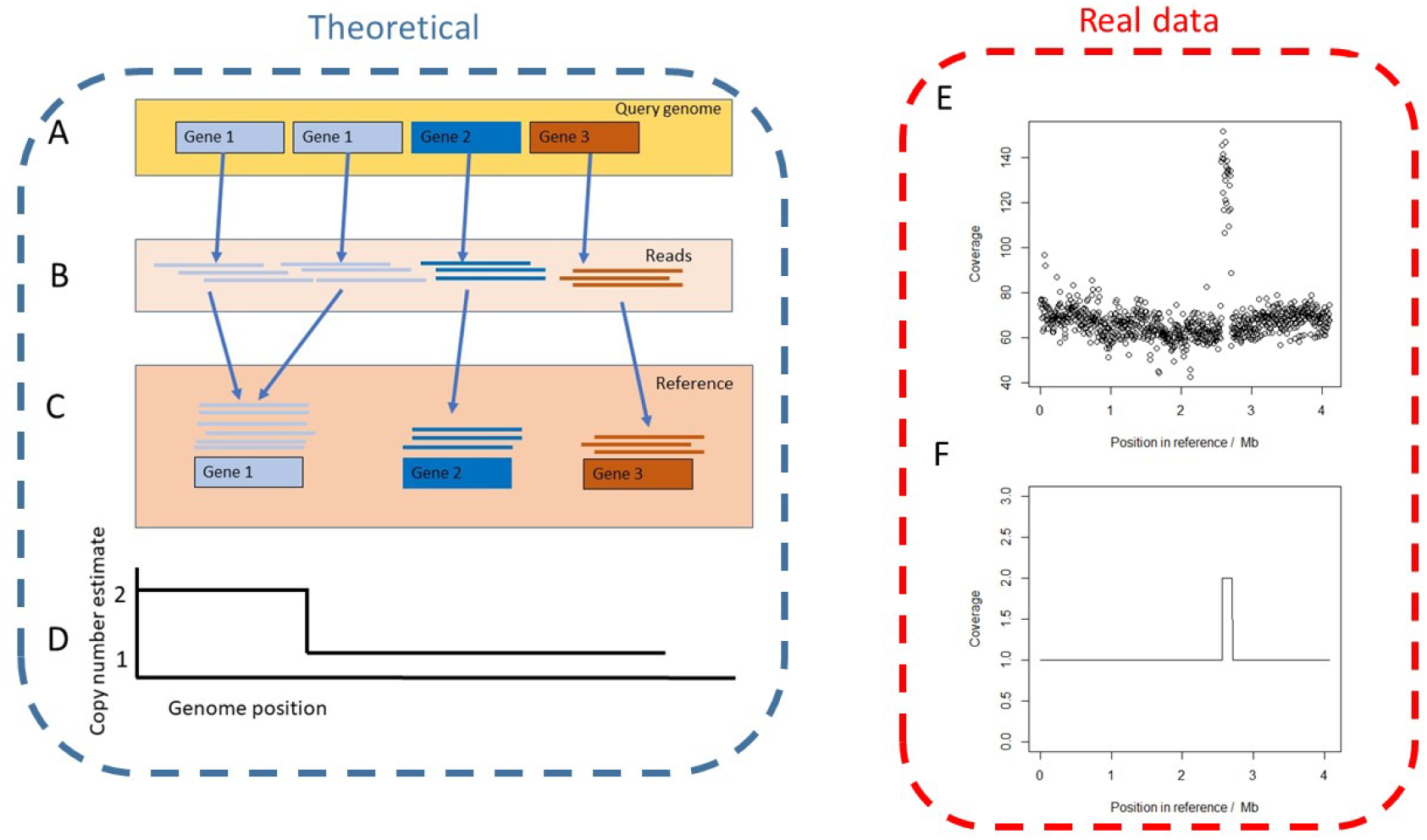
Schematic overview of prediction of CNVs from sequencing read depth. In the theoretical example (purple box, left), the query strain contains a perfect tandem duplication of gene 1 whilst gene 2 and 3 are at single copy (A). Short reads from the query strain are generated (B) and mapped to the reference genome, that contains all genes at single copy (C). Reads from both copies of gene 1 in the query strain map to this locus in the reference sequence and thus twice as many reads map to this gene compared to genes 2 and 3. This data must be processed to avoid technical bias, the pipeline processes read coverage data into estimates of copy number (D). Using an example with real data (red box, right) the strain SAMN08200079 was analysed. Read coverage was graphed to reveal a duplication at ~1.4Mb (E, analogous to theoretical graph C) which was statistically analysed using our pipeline (F, analogous to theoretical graph D).

The performance of our approach was first tested using Illumina HiSeq reads simulated from the B1917 reference genome, which does not contain CNVs. As expected, no CNVs (false positives) were predicted and all genes were correctly estimated at single copy. The approach was further evaluated by mapping Illumina data from those strains with manually resolved CNVs described above, each of which contained one CNV.

When data is mapped to a reference the true gene order of the sample is masked- an inherent feature of read mapping. Therefore, strains with duplications in rearranged loci may appear as discontinuous stretches of duplicated DNA in the reference. When establishing the accuracy of the pipeline we only considered resolved CNVs that were contiguous on the B1917 reference genome as other CNVs are impossible to accurately resolve. Two samples were therefore excluded because they contained rearrangements relative to B1917. Whilst the 25 remaining CNVs occurred at just three distinct loci, their beginning and ending coordinates, as well as overall length, varied between strains. Thus, three measures of accuracy were tested: the correct prediction of the 25 CNVs, the quantity of false positives and the predicted beginning and ending locations of each CNV (breakpoint accuracy). Only one (J321) of the 25 data sets failed our quality control (see Methods) for high read depth noise and was excluded; leaving 24 high quality strains.

Of the 24 resolved, high quality and suitable CNVs, 23 were correctly predicted (defined as >=80% reciprocal overlap) (Supplemental_Table_1). Three false positives were detected in three different strains. Two of these were due to one gene within the CNV locus being predicted as single copy, causing the true, single CNV to be predicted as two, separated by the falsely predicted single copy gene. In the third false positive, a second locus was predicted as a duplication and despite further analysis, no evidence was found of a second duplication in this isolate.

The breakpoint accuracy of estimates was calculated with false positives excluded (Supplemental_Figure_1). The median distance between the true values and the read depth-based estimates was 1 gene. There were five estimated start/end points which were considerably (>=5 genes) less accurate than the rest of the dataset, mainly arising from the two strains in which the CNV was predicted as two separate loci.

Thus the pipeline correctly predicted, and with excellent breakpoint accuracy, the CNVs for 20 of the 27 resolved genomes (74%), with 3 further CNVs predicted (11%) but as two adjacent but separate loci.

### CNVs as a source of genetic diversity

The pipeline was applied to predict CNVs in 2709 *B. pertussis* isolates for which short-read sequence data was available in the Sequence Read Archive (SRA) or locally provided (n=94). Of the 2709 total *B. pertussis* samples, 94 exhibited < 30x average coverage and 185 had high read coverage noise. Therefore, the final test dataset included 2430 *B. pertussis* isolates (Supplemental_Table_2). B1917 was used as the reference genome. Of the 2430 studied isolates, 1711 had all genes predicted at single copy, leaving 719 strains with at least one deletion or CNV. Of these, 191 isolates contained 272 CNVs- some strains containing multiple CNVs. Computed copy number estimates (Supplemental_Table_2) were visualized with an interactive heatmap for inspection where it became apparent that particular loci were present as CNVs in multiple strains, which we termed ‘hotspot’ loci (Figure 2). Consistent with our observations in the resolved dataset and previous reports (Ring et al. 2018; Weigand et al. 2016, 2018a), CNVs at hotspot loci varied in length between isolates, with differing start and end points but including a core set of genes.

**Figure 2.**
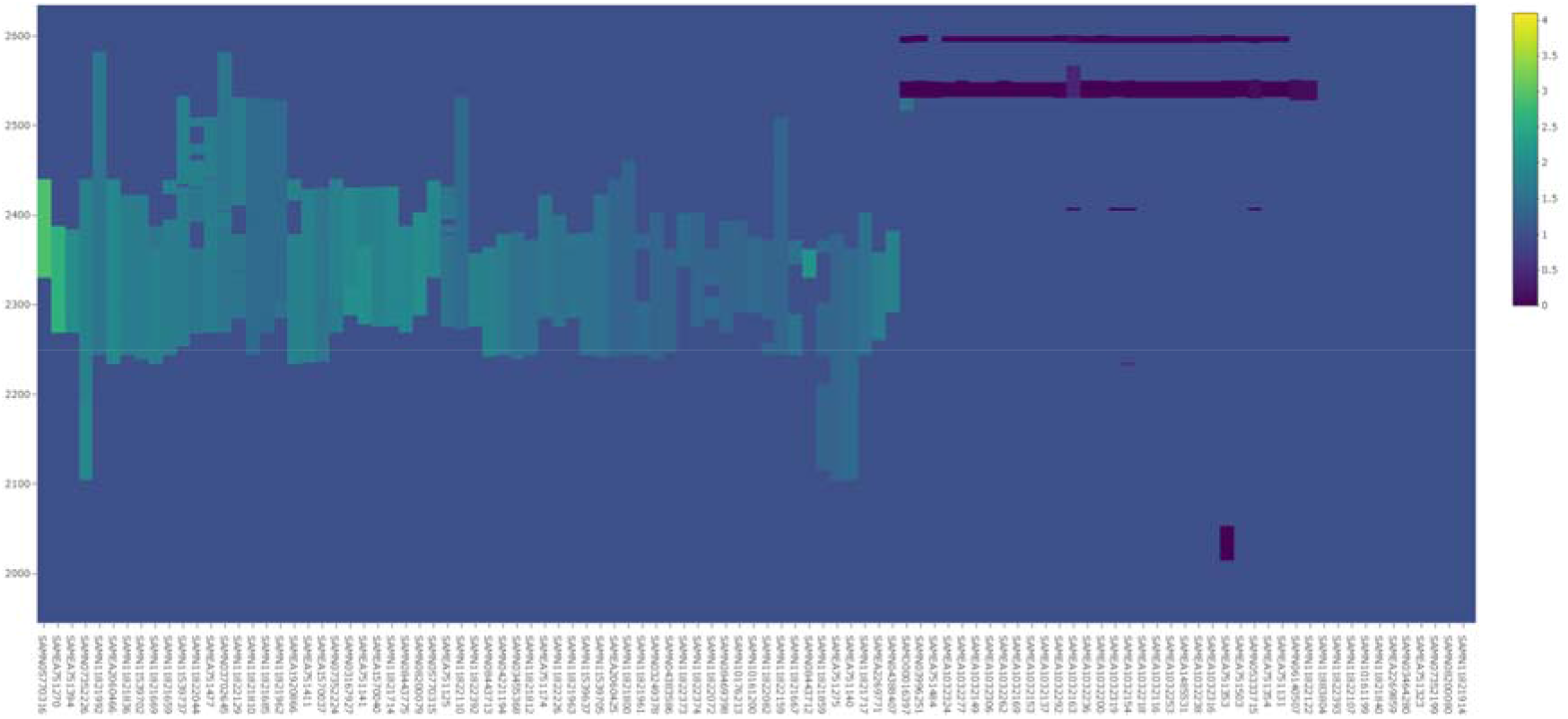
A section of the heatmap comprising the majority of CNVs in Network 1, including a triplication of this locus. Isolates are in columns, while rows indicated the index of each gene in the reference sequence. The colour scale (Z axis) indicates the copy number of each gene. A legend of the colour scale is on the far right.

### Most CNVs occur at hotspots

The relationship between all CNVs was quantified as the proportion of gene content overlap between all pairwise comparisons. Network graphs were constructed between CNVs (‘nodes’) that were connected by at least 75% content overlap (‘edges’). The 272 identified CNVs formed 24 network graphs, representing 24 distinct genomic loci (Supplemental_Table_3). Only 11 network graphs, corresponding to the hotspot loci, included three or more isolates and contained 254/272 (93%) of the predicted CNVs (Supplemental_Figure_3). Network density is the percentage of theoretically possible edges observed between nodes in a network. The mean density of all CNV networks was 71%, indicating that the CNVs in each network were highly interconnected, Table 1. https://plot.ly/~kows1337676/433.embed

**Table 1.** ‘Hotspot’ CNV network statistics.

| Network name | Frequency (CNVs) | Mean length (genes) | Median start (B1917 gene name) | Median end (B1917 gene name) | Mean copy number | Core (>=55% proportion (%)) | Network density (%) |
| --- | --- | --- | --- | --- | --- | --- | --- |
| 1 | 102 | 106 | RS12140 | RS12755 | 1.6 | 50 | 55 |
| 2 | 57 | 82 | RS15100 | RS15490 | 1.7 | 61 | 63 |
| 3 | 21 | 80 | RS07175 | RS07660 | 1.68 | 35 | 60 |
| 4 | 18 | 20 | RS00010 | RS00130 | 1.35 | 96 | 100 |
| 5 | 13 | 67 | RS19230 | RS19625 | 1.93 | 51 | 50 |
| 6 | 11 | 75 | RS05505 | RS05935 | 1.6 | 77 | 73 |
| 7 | 8 | 49 | RS04185 | RS04430 | 1.88 | 78 | 71 |
| 8 | 8 | 74 | RS09665 | RS10290 | 1.82 | 58 | 43 |
| 9 | 7 | 13 | RS19965 | RS10580 | 2.49 | 98 | 100 |
| 10 | 6 | 23 | RS19465 | RS19565 | 1.32 | 94 | 100 |
| 11 | 3 | 45 | RS01035 | RS01300 | 1.63 | 87 | 67 |

Given the density of CNV networks, hotspot loci were investigated further to identify which genes formed the ‘core’ and how central these genes were to their respective network. To define the network core, we determined the genes contained in at least 55% of the CNVs in a network. To extend beyond just defining the central genes of the network we quantified the difference between the central core and the full-length CNVs within each network (analogous to the core and accessory genome in the study of pangenomes). The mean overlap between the CNVs in a network and the network core was calculated to give the ‘core representation’ statistic. These metrics revealed that the core of most networks, with the exception of network 3, comprised at least 50% of the mean length of all CNVs in each network (Supplemental_Table_4). Thus, the 11 hotspot loci described here were composed of CNVs that varied around a central core rather than overlapping CNVs arranged in series.

It could be seen in the heatmap that not only did the CNVs cluster at specific hotspots but that some samples had multiple CNVs at the same hotspot. This may have been due to a complex mixture of structural variations affecting the same locus, such as nested duplications (Weigand et al. 2018a) or the locus being disrupted by inversions. It is also possible that, as detected in two cases of the benchmarking experiment, a CNV was predicted as two separate regions of higher copy number. A number of network cores contained genes with varied, predicted functions. For example, Network 1 contained genes for flagellar motility (Hoffman et al. 2019); Network 2 contained the *nuo* operon which is linked to respiration (Nakamura et al. 2006; Archer and Elliott 1995) and Network 3 contained the *fim3* gene involved in the pathogenesis of *B. pertussis* and present in some acellular vaccine formulations (Scheller et al. 2015). In addition, the networks contextualised the 25 resolved genomes previously studied, the majority of which were in networks 1, 2, and 3.

### CNV plasticity during *in vitro* growth

CNVs identified above were often predicted with non-integer copy numbers in addition to copy number discrepancies between predicted and resolved copy number in the manually resolved dataset (Supplemental_Figure_2). To confirm our predictions from short-read sequencing data and investigate the basis for non-integer copy numbers, we exploited the tractable size of one relatively small CNV. The genome of UK54 (SAMEA1920853) was predicted to have a 16 kb CNV at a copy number of 4.1; short enough to observe the entire CNV locus in a single sequence read on the Nanopore platform, assuming that each copy occurred in tandem as observed in both our data and previous reports (Weigand et al. 2016, 2018c). The duplication was part of Network 9 which was comprised of 7 other duplications, one of which was also predicted at a copy number >2 (3.3, Strain SAMN11822098).

The copy number of this locus in UK54 was first validated using qPCR. The relative copy number of a gene within the CNV compared to a single-copy gene encoded outside the CNV locus was 4.38 +/− 0.4 which matched the read depth-based prediction.

Whole genome sequencing on the Nanopore platform yielded a mean read length of only 9.1kb but produced over 3000 reads with a length exceeding 50kb. Sequence reads that contained both of the regions flanking the CNV locus and the CNV locus itself were identified (n = 9) and contained the CNV at different copy numbers (Figure 3). This demonstrated that a laboratory culture of UK54 comprised a mixture of copy numbers at this locus and explains the non-integer copy numbers predicted by CNVnator. Genomic DNA for sequencing is derived from laboratory populations of bacteria and if these harbour CNVs at different copy numbers, subsequent read-depth based predictions will represent the average read depth of all of the bacteria sequenced.

**Figure 3.**
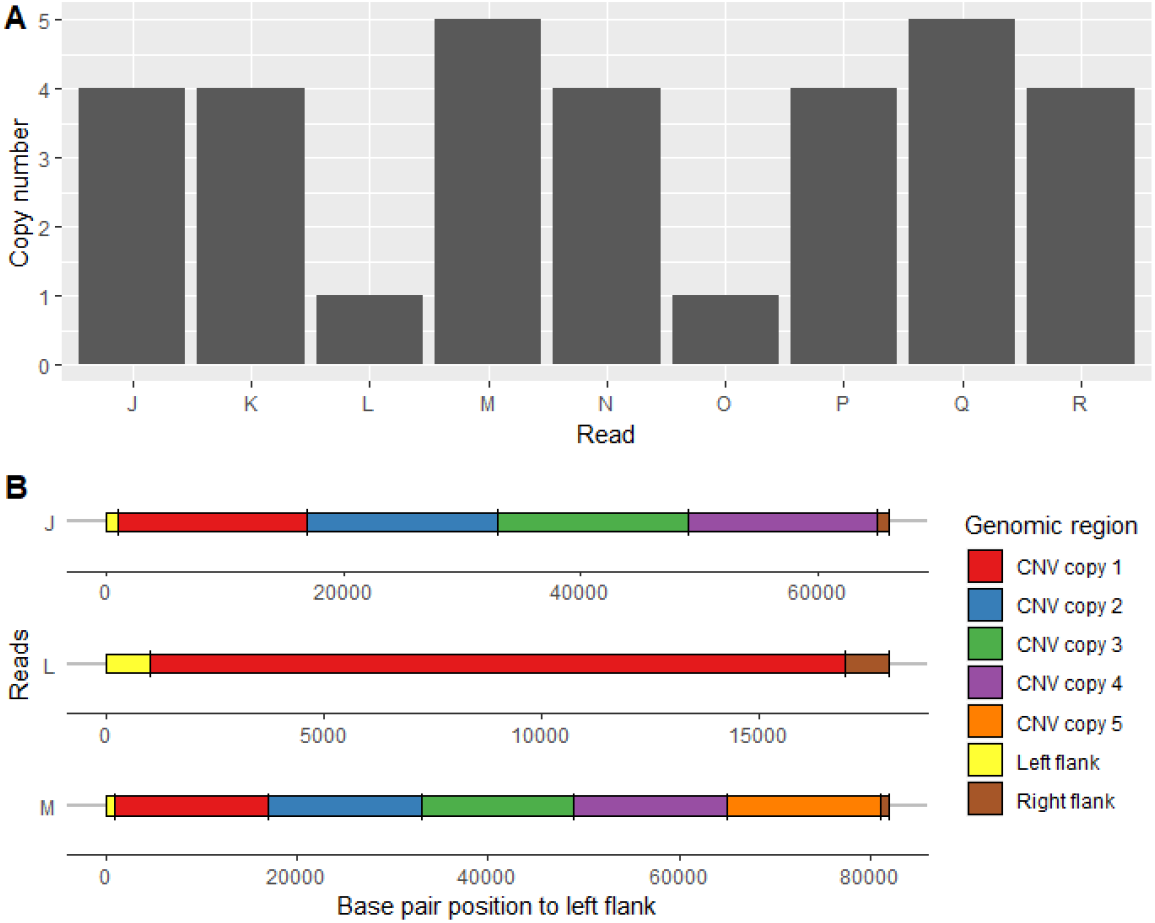
Ultra-long read sequencing of UK54 revealed the presence of different copy number CNV loci within a single culture. Individual sequence reads that spanned the CNV loci were identified using Blastn, labelled J to R. (Panel A). The data shows each read (x-axis) containing 1,4 or 5 copies of the locus (y-axis) and therefore, as each read appears to be integrated into the chromosome, there were cells present in the population with 1, 4 or 5 copies of the locus. The arrangement of the relevant section of three reads (J, L and M) is illustrated in panel B.

It was not known if the original culture of UK54 involved isolation of a single colony or collection of multiple clones from the diagnostic plate growth and thus whether the observed variation in copy number resulted from a mixed culture or emerged during laboratory growth prior to sequencing. To investigate this, we picked eight single colonies of UK54 and passaged them by growth on agar and then during broth growth. Each of these clonal populations were theoretically derived from a single bacterium. The copy number at the CNV locus in each of the resulting clones was estimated using qPCR (Figure 4) and ranged from 2.2 (clone 6) to 51.2 (clone 8). We sequenced UK54 clone 4 using the Nanopore platform and observed sequence reads with copy numbers 1, 2, 4, and 5 (Figure 5). These data strongly suggested that CNV copy number was plastic, with variants arising during *in vitro* growth from a single bacterium to the culture from which the gDNA was extracted.

**Figure 4:**
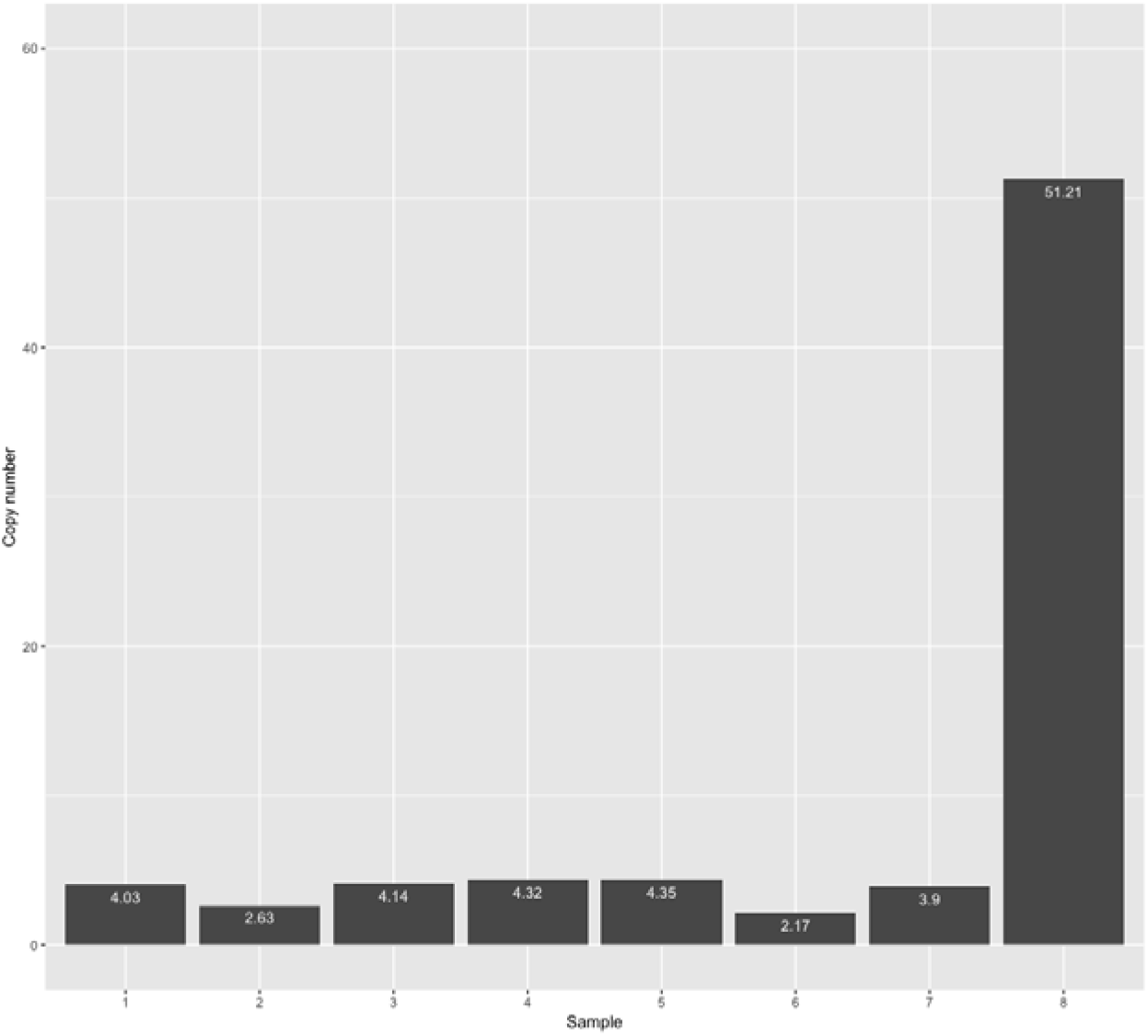
Quantification of CNV copy number of 8 clones of UK54 by qPCR demonstrated a range of copy numbers from 2.17 to 51.21.

**Figure 5.**
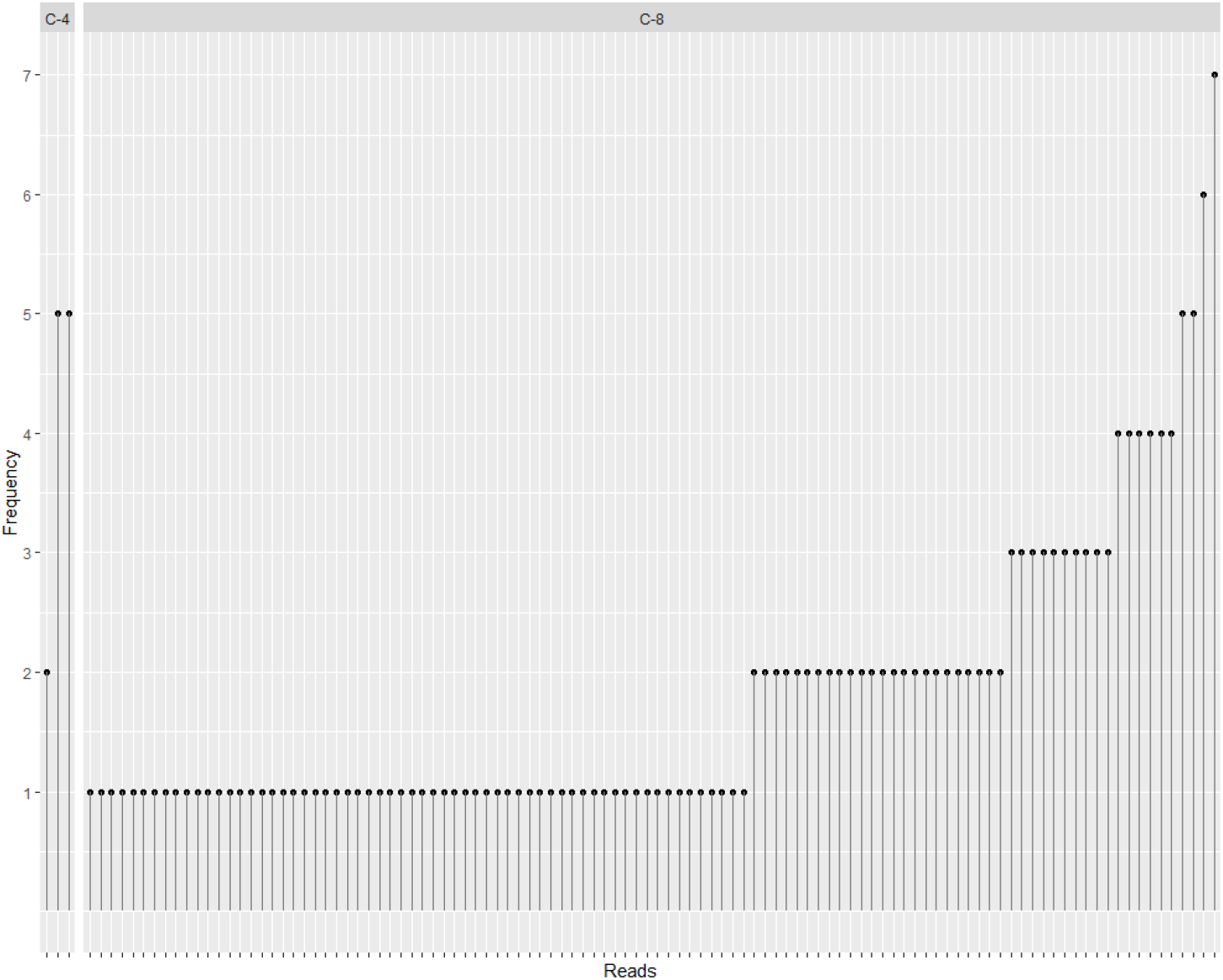
Nanopore sequencing of UK54 clone 4 and 8 (C-4 and C-8) revealed the presence of different copy number loci within a single culture. Individual sequence reads that spanned the CNV loci were identified using Blastn successfully in clone 4 whilst clone 8 had no reads spanning the full locus. The data show each read (X axis) contained between one and seven copies of the locus (Y axis).

Nanopore sequencing was also performed with UK54 clone 8, which exhibited a copy number estimate of 51 by qPCR (corresponding to a predicted CNV length of 816kb) (Figure 5). No reads spanning the entire CNV locus (i.e. the CNV locus with flanking DNA on each side) were produced, presumably due to its extreme length. However, reads containing up to 7 copies of the locus, without flanking regions, were identified. Relaxing the Blastn alignment parameters from a 90% minimum query length of the CNV locus to 50% identified a maximum of 9 copies of the locus present on a single read with incompletely sequenced copies at each end. Consistent with the copy number prediction from qPCR, the read depth at this locus for UK54 clone 8 from the Nanopore data was approximately 60x higher than the genome average, strongly supporting the very high copy number estimate for this locus in this clone.

To investigate potential phenotypic variation resulting from amplification of genes by CNV formation, we measured mRNA levels for one gene within the CNV locus in UK54 clones with different average copy numbers. Levels were normalized to the single copy *recA* that is often used as a stably-expressed housekeeping gene in RT-qPCR experiments. We selected clones 2, 4, and 8, with screened copy numbers of 2.63, 4.32, and 51.21, respectively. As we demonstrated that each culture comprises a heterogenous mixture of cells with varied CNV copy number, we re-estimated the locus copy number for each clone using the same laboratory culture from which RNA was extracted. Upon re-growing these clones for RNA extraction, the average copy number in each changed (non-significantly) to 4.1, 6.5, and 53.1 in clones 2, 4, and 8, respectively. The mRNA level for the CNV gene corresponded with the copy number (Figure 6); normalising the transcript level in clone 2 to a value of 1, it was 16.8 fold higher (P<0.0001) in clone 8. It was also higher, but not significantly, in clone 4 (P=0.76). However, broadly, using the data as a whole, there is an association between DNA copy number and transcript abundance. This strongly suggests that the gene dosages produced by CNVs affected relative gene expression levels.

**Figure 6:**
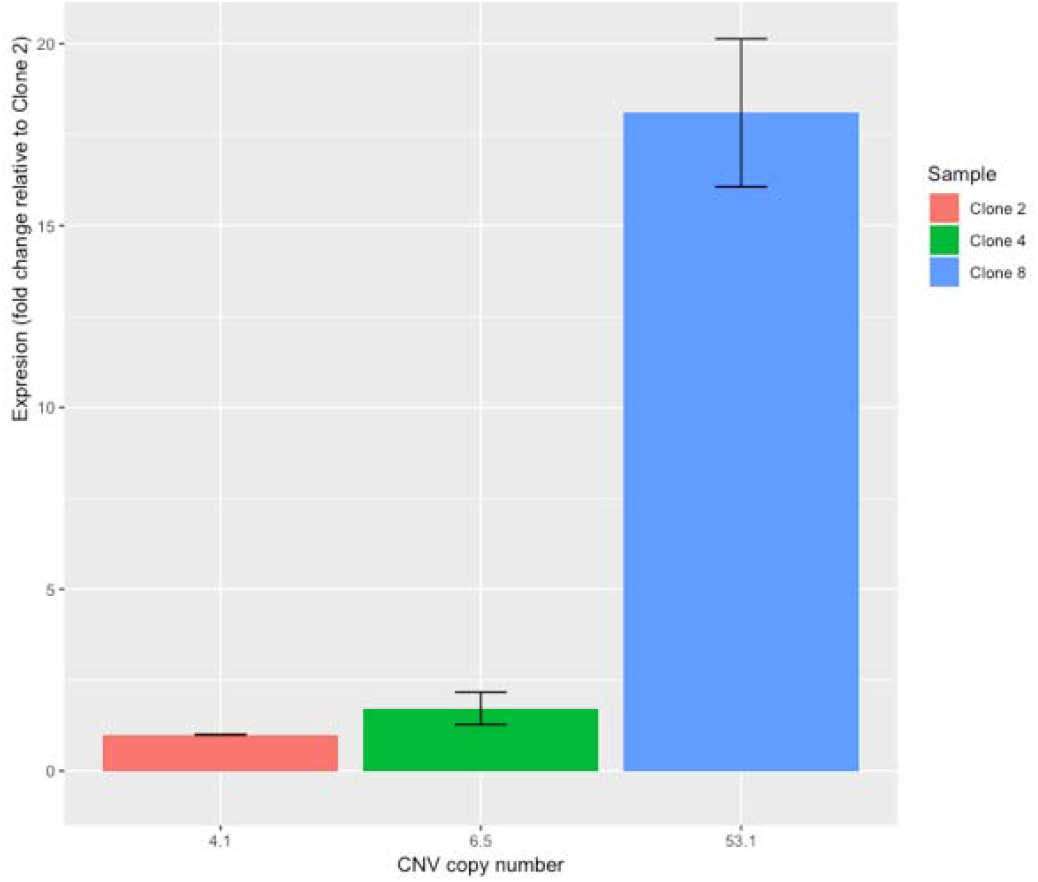
Copy numbers of clones 2, 4 and 8 were quantified using qPCR and expression of a gene within the CNVs was quantified by RT-qPCR. Expression is shown as a relative fold change to Clone 2. Error bars represent standard deviation of expression. The results show that copy number corresponds to RNA expression.

### Structural plasticity during *in vitro* growth

Analysing the Nanopore data from clonally derived populations strongly suggested that the CNV locus in UK54 was plastic. To investigate if similar effects were occurring at other genomic loci we identified sequence reads that contained regions that are proximal on the sequence read but not in the consensus genome sequence-putative structural variants arising from genome rearrangement. In UK54 clone 4, 59 reads out of >600k total reads were identified as having a gene order that was different to the consensus sequence. Structural variants occur primarily by recombination between repeats (although recombination is possible between regions with no homology (Reams and Neidle 2004; Nilsson et al. 2006)) and therefore, only the 22 reads containing repeat sequences at the junction were further analysed.

Structural variations could be delineated between rearrangements or deletions/CNVs by analysis of the orientation of the DNA before and after the junction. If the DNA segments are in the opposite orientation it can be assumed the mutation was an inversion whereas if they are the same orientation it is likely the mutation was a deletion or duplication as has been seen previously (Weigand et al. 2017, 2018b).

Our results demonstrated that, like CNV copy number, putative structural variants can also be detected during *in vitro* growth, distributed around the genome (Figure 7). Interestingly, in addition to structural variation via recombination between IS, we observed both rearrangement and deletion/CNV from recombination between a 3kb locus found duplicated in a number of recent clinical isolates and these duplicated loci were identified as a potential hotspot for recombination (Figure 7) (Weigand et al. 2017). Tandem CNVs between these sequences were not observed in our study of 1000’s of isolates but Weigand et al found that rearrangements arising from recombination between these loci were common (Weigand et al. 2017).

**Figure 7.**
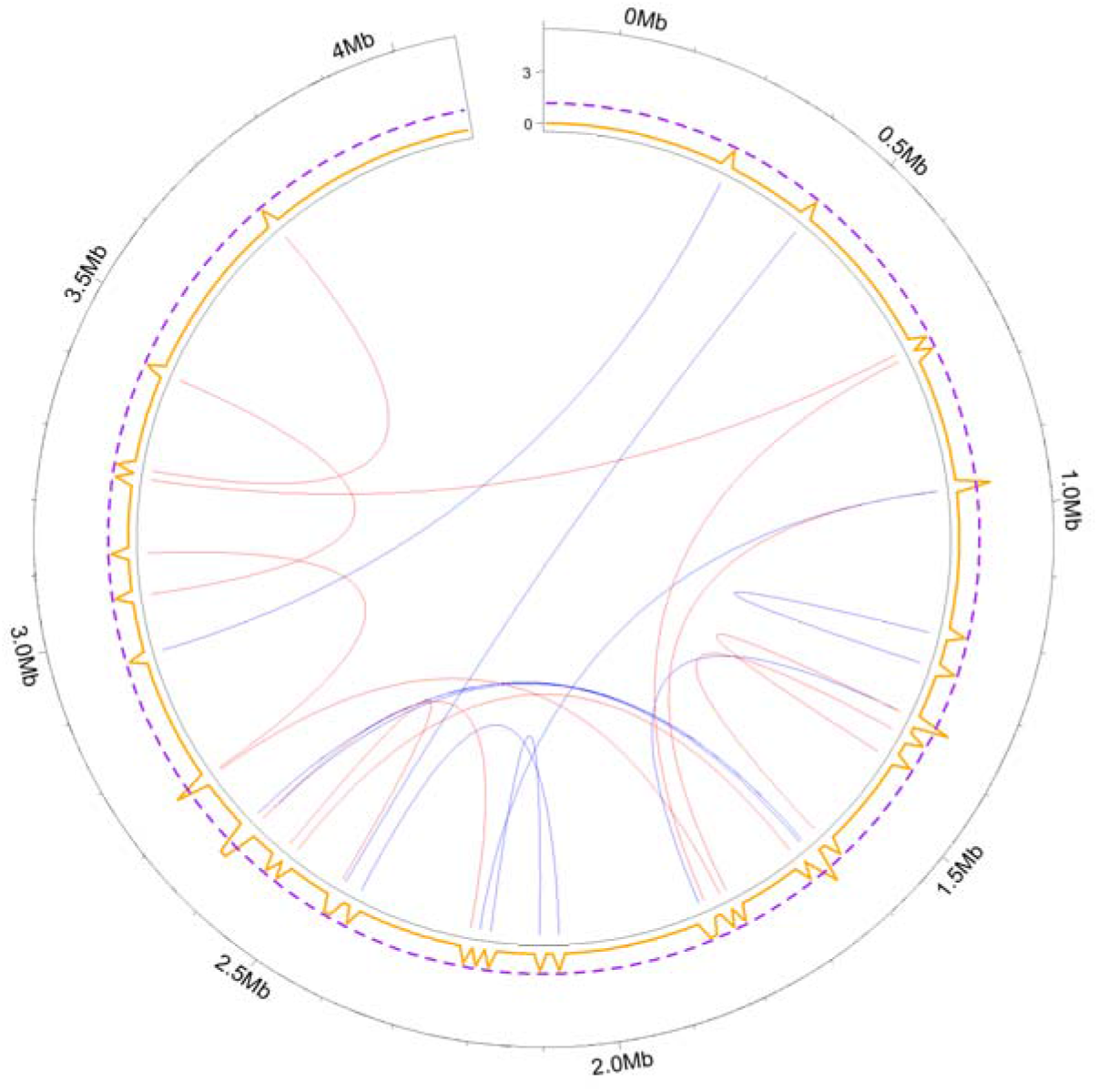
Circos plot (inner circle) of putative structural variants present in individual reads (n=22) that were absent in the consensus sequence. Red lines are CNVs or deletions and blue lines are rearrangements. The frequency of structural mutations occurring in 10kb windows was plotted (orange line) with the dashed purple line representing three standard deviations above the mean to identify potential ‘hotspot’ regions. The analysis shows a uniform distribution of structural variants around the genome with only the regions at 1.6Mb and 2.6Mb representing a potential hotspot.

Often, clonal bacterial cultures are sequenced to study mutations that have occurred further back in evolutionary history and have become fixed. However, these experiments also inadvertently capture a ‘snapshot’ of evolutionary time. We can therefore observe the creation of a variety of errors in the DNA of populations of cells using this ‘snapshot’ – mutations which are otherwise invisible when an average (consensus) sequence is made for the population. Whilst cells with lethal or highly deleterious mutations are not expected to persist in a population, a number of reads appeared to strikingly indicate deletions or duplications of over 1Mb of DNA. Whilst these structural variants are putative, they indicate, in combination with our other results, the ongoing genome plasticity of BP.

### CNVs were highly associated with repetitive sequences

While we verified one predicted CNV, verification was not feasible for the remaining 272. However, to increase confidence in these predictions we investigated their association with repetitive elements, compared to all genes. All previously published and resolved CNVs were adjacent to repetitive sequences, suggesting this was a clear marker for true CNVs. We used only closed genome sequences, as the location and frequency of repetitive sequences varies between strains, and excluded CNVs already described in the manually resolved genomes or which were disrupted by genome rearrangements, leaving 16 CNVs in 13 isolates.

The 16 predicted CNV boundaries were significantly (p<6^−08^) closer to repeat genes (median distance of +/− 1 gene) than non-CNV genes (median distance of +/− 5 genes) (Figure 8 and Supplemental_Table_5). This, in conjunction with our stringent quality control steps and the previously accurate predictions, supports the accuracy of the prediction of 272 CNVs.

**Figure 8.**
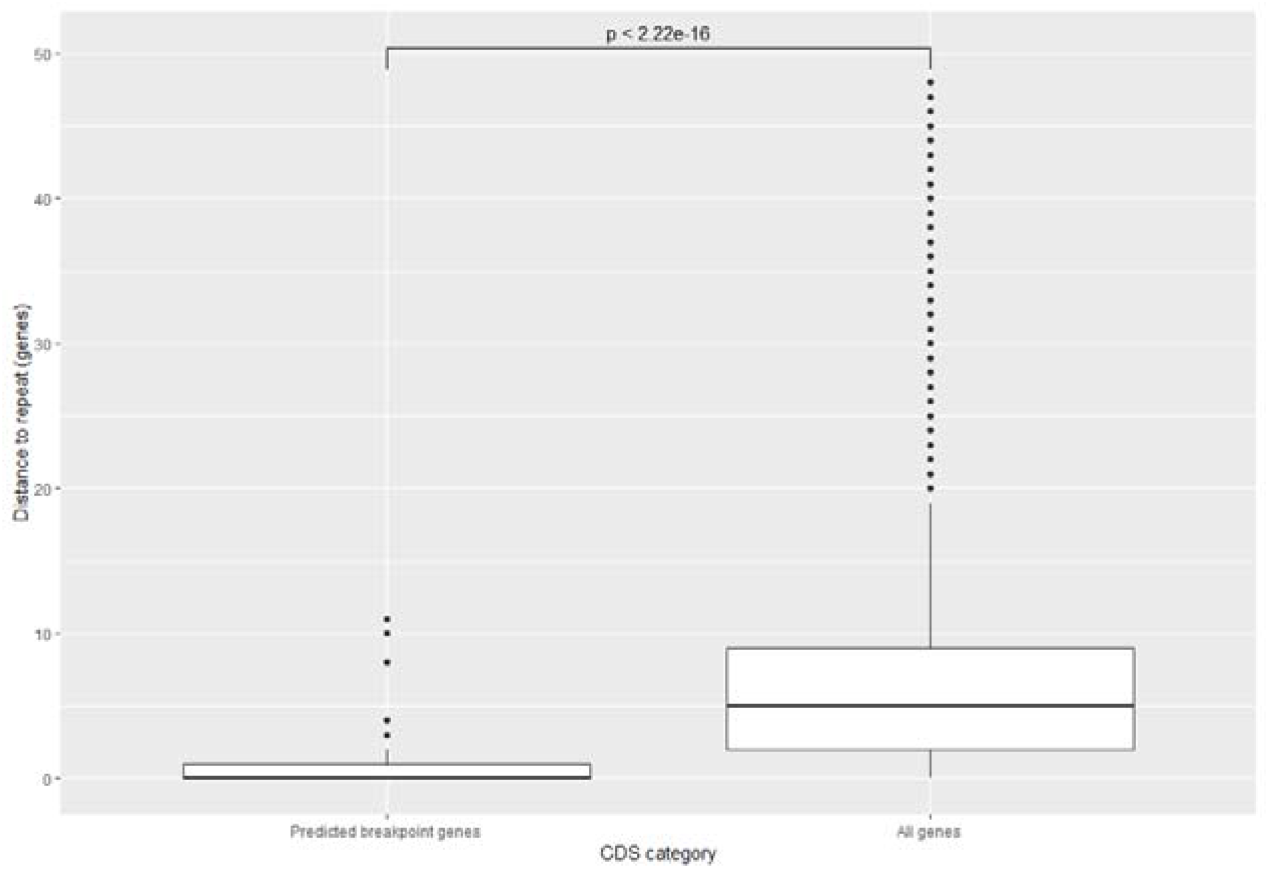
The distance (measured in genes) between CNVs and repeat genes was identified in closed genomes. The ends of CNV loci were found to be significantly closer (median: 0 genes) to repeats than the average gene (median: 5 genes).

### CNVs occur sporadically throughout the phylogenetic tree

We had demonstrated that CNVs could change copy number over the microevolutionary timescales of days during growth in the laboratory. In addition, it was demonstrated that while CNVs did overlap at hotspot loci they often had unique gene contents-strongly indicating each arose from an independent mutation. Therefore, we theorised that there was no strong phylogenetic relationship between CNVs within a network. To test this, we sought to estimate for each network if at any point in the phylogenetic tree an ancestral strain, represented by an internal node of the tree, was likely to have had the corresponding CNV (Supplement_Table_6). This was performed by using presence or absence of CNVs within a network as a discrete trait.

Ancestral state reconstruction (ASR) resulted in 16 nodes near the tips of the tree (<=7 SNPs and <=8 tree splits) having a high likelihood (>0.8 empirical Bayesian posterior probability) of being in a duplicated state. Due to the large number of isolates studied in combination with the extremely low diversity of *B. pertussis*, however, branch lengths were often 0-potentially skewing the results. Further investigation of this effect showed that 7 of the 16 nodes of interest had just one tip with branch length 0 directly stemming from the node. The extremely close relationships between these tips and nodes leads to an overwhelming statistical signal to the ASR algorithm leading to false positive results. These 7 nodes were ignored. Our results therefore indicated very limited heritability of CNVs, but yielded 8 putative examples of the mutation being maintained over small evolutionary time scales.

## Discussion

*B. pertussis* is described as a monomorphic bacterium (Mooi 2010) that has evolved as a human-specific pathogen through gene loss via homologous recombination between direct repeats. However, homologous recombination can also cause multi-gene CNVs. Although 12 multi-gene CNVs had been described previously, no systematic analysis of CNVs in *B. pertussis* had been carried out. In this study, short-read genome sequence data generated on the Illumina platform for 2430 strains were analysed using read depth as a proxy for copy number. Our results revealed 11 clusters consisting of 272 CNVs, some of which comprised hundreds of genes, revealing a novel aspect of genetic variation among *B. pertussis*. This contributes to a growing literature that demonstrates that quantifying *B. pertussis* diversity requires a comprehensive view of mutation types, not just the quantification of DNA base changes (Weigand et al. 2017; Bowden et al. 2016; Weigand et al. 2018a).

The large number of copies of IS*481* throughout the *B. pertussis* genome suggests that a very large number of different genome rearrangements (Weigand et al. 2017) and CNVs are possible. Despite the vast diversity of possible CNVs, however, 94% of observed CNVs appeared at just 11 hotspot loci, suggesting strong purifying selection acts on CNVs in *B. pertussis*. This discrepancy between the potential and observed distribution was further explored by sequencing strains after limited *in vitro* growth – greatly reducing (but not eliminating) the effects of selection. We also identified putative *de novo* generation of structural variants, albeit infrequent, and demonstrated that the copy number of the studied CNV (in UK54) was plastic over short laboratory timescales. High genome plasticity was further supported by the limited heritability identified using ancestral state reconstruction among the global *B. pertussis* population. Our results shed light on the continual homologous recombination in *B. pertussis* and support a range of studies that established the genome dynamics of homologous recombination in bacteria (Anderson and Roth 1981; Edlund et al. 1979; Chen et al. 2008).

CNVs can be very costly mutations, carrying as much as a 0.15% fitness cost per 1kb, primarily due to increased gene dosage leading to additional transcription and translation rather than replication of a larger genome (Adler et al. 2014). According to this estimate, the larger CNVs observed in this study may carry fitness costs over 30%. Therefore, unless higher levels of transcribed (non-coding RNA) or translated (proteins) gene products provided a strong selective advantage to overcome such a cost, the CNVs is likely to be selected against. One of the most frequent hotspots observed in this study included genes for flagellar motility. Motility has been frequently implicated in the virulence of bacterial pathogens, but *B. pertussis* has long been regarded as non-motile. However, recent research has shown that motility can be occasionally observed *in vitro* (Hoffman et al. 2019) and flagellar biosynthesis genes are expressed during murine challenge compared to *in vitro* growth, potentially implicating motility or biofilm formation in infection (van Beek et al. 2018). Duplication at this locus may, therefore, affect the virulence, colonisation, or carriage of *B. pertussis* in the human population, but the influence may be modulated due to the plasticity of CNVs.

Long-read Nanopore sequencing of two clones of UK54 led to the remarkable observation of a CNV with average copy number of 51. It is a well-documented phenomenon that multiple copies of a locus in tandem greatly increases the instability of the locus. In experimental systems, copy numbers of up to 100 have been generated (Edlund et al. 1979) and in clinically derived isolates the copy numbers of antimicrobial resistance genes can change rapidly, increasing up to 70 copies in response to antibiotics (Nicoloff et al. 2019). Whilst the function of the genes in the UK54 CNVs are unclear it is possible that they provide a fitness benefit to the strain under certain conditions.

Our investigation demonstrates several widely applicable approaches to the study of CNVs. Our application of a CNVnator-based pipeline utilises short-read sequencing data that is available for thousands of bacteria. A limitation of this approach is that reads were mapped to B1917 and therefore CNVs were predicted as if the gene order was the same in B1917, despite frequent rearrangements in the population. This may lead to the read depth signal being ‘split’ on the B1917 reference when they are, in truth, contiguous on another genome, leading to multiple CNV predictions. To overcome the limitations of short read sequencing we therefore used long-read DNA sequencing for spanning repeat regions to enable resolution of CNVs and genome arrangement, although correct assemblies required additional data. However, genotypes could be described using single Nanopore reads to identify copy number heterogeneity within in-vitro cultures.

Network graphs were used to analyse the complex relationships between CNVs in *B. pertussis* quantitatively. This arrangement of CNVs appears in other bacterial species (Weiner et al. 2012) in addition to plants (Faris et al. 2000) and animals (Perry et al. 2006) and therefore, networks are a flexible and generic framework to analyse such phenomena. An advantage of using networks to describe hotspot loci was the ability to semantically categorise CNVs to unite the findings of many studies and contextualise them with new data. Previously, using limited data, we had demonstrated a ‘hotspot-like’ effect by resolving four CNVs with subtle gene content variations at the same loci (Weigand et al. 2016, 2018a), corresponding here to Network 1, at which other CNVs had also been reported (Dalet et al. 2004; Weigand et al. 2018b, 2016; Heikkinen et al. 2007). Our results contextualise this research, providing another 90 CNVs at this location and we combined core CNV and mean overlap statistics to show that the majority of the 11 networks in our analysis consisted of CNVs which varied around a core set of genes, just as in Network 1. The varied start and stop positions of overlapping CNVs among strains offers further evidence that amplification of the core genes may be under selection yielding multiple independent mutations.

The unusually high number of insertion sequences within the *B. pertussis* genome, and their relatively even distribution, likely facilitates the genome-wide distribution of structural variants. Indeed, genomes of related species *B. parapertussis* and *B. holmesii* each harbour fewer IS elements and thus exhibit fewer rearrangements (Weigand et al. 2019) and very rare CNVs (M.R. Weigand, unpublished). However whilst unusual, *B. pertussis* is not unique, as its abundance of IS elements ranks in the top 30 in a study of 1000’s of bacterial isolates (Robinson et al. 2012). It is likely that such dynamics are playing out in other species proportional to their IS elements load (Weiner et al. 2012; Yang et al. 2005).

In conclusion, we have rigorously and successfully investigated the repertoire of CNVs in *B. pertussis*, revealing a novel layer of diversity that should be considered when quantifying variation within the species. These results revise existing knowledge of circulating *B. pertussis* and highlight challenges to molecular surveillance. Previously, low-resolution genome typing, specifically pulsed-field gene electrophoresis (PFGE), has been the primary tool for pertussis molecular epidemiology due the high diversity of profiles observed among clinical isolates compared to other methods (Bowden et al. 2014). While much of this diversity is attributable to rearrangement (Weigand et al. 2017), the present study also highlights a role for CNVs and that their transient and homoplasic nature may mask the ancestral (epidemiological) relationships between strains or over-estimate genetic diversity. (Weigand et al. 2017) Taken together, our contemporary genomic study of circulating *B. pertussis* should signal the end to this pathogen’s designation as a monomorphic species.

## Methods

### Sequence read mapping

Short read data originating from the Illumina platform were retrieved from the National Centre for Biotechnology Information’s (NCBI) Sequence Read Archive (SRA). One run was chosen at random for each BioSample, totalling 2709 runs including 94 locally provided runs. Reads were mapped to the *B. pertussis* B1917 genome, which is broadly representative of the modern circulating strains (Bart et al. 2014) (RefSeq ID: NZ_CP009751.1), using BWA (Li 2014) implemented in Snippy (available: https://github.com/tseemann/snippy).

### CNV prediction

CNVnator (Abyzov et al. 2011) was used to predict CNVs from read depth data generated from the mapping process. Statistical tests for significance within CNVnator discriminate high and low confidence calls. To further increase specificity, we implemented a very low P-value cutoff (*p*<0.0001). Abyzov *et al* empirically tested CNVnator to determine that ratios of the average read depth to the standard deviation of 4-5 produce the best balance between sensitivity and specificity (Abyzov et al. 2011). In accordance, samples exhibiting ratios < 3 were discarded as CNV calls were unreliable on such variable data (Abyzov et al. 2011). Window length was optimised for each genome, testing window sizes 500-1000bp at intervals of 100bp to evaluate which gave a ratio closest to 4.5 as to minimize the effect of stochastic and/or artefactual fluctuations in read depth across the genome. Copy number estimates were rounded to the nearest 0.1. Code is available: https://github.com/Jonathan-Abrahams/Duplications

### Control data

As a negative control, short reads were simulated from the B1917 reference genome using ART to simulate the error profile of Illumina HiSeq paired-end 150 bp data (-ss HS25 -p -| 150 -f 20 -m 200 -s 10) (Huang et al. 2012). Simulated reads were mapped back to the reference genome using Snippy and CNVnator was used to call any spurious CNVs, as described above (Abyzov et al. 2011). As a positive control dataset, closed genome sequences from 25 isolates with manually resolved CNVs were used. This data was generated using a combination of PacBio and Illumina sequencing and optical mapping on the Argus or Nabsys HD platforms, as done previously (Weigand et al. 2016, 2017, 2018c).

### Heatmap

The read depth-based predictions were hierarchically clustered based on the similarities of CNV profiles (including deletions) of samples using the R package Hclust. This therefore meant that strains with similar complements of CNVs and deletions were clustered together on the heatmap which was plotted using the R package Plotly (Plotly Technologies Inc. 2015).

### Networks

Overlapping gene content among CNVs was evaluated by constructing undirected network graphs which quantified the relationships (edges) between each CNV (nodes). An edge was constructed between nodes if both CNVs had a 75% overlap (non-reciprocal). Network analysis was undertaken in R using the Igraph package (Csardi and Nepusz 2006) and networks layout was generated by the Fruchterman algorithm (Fruchterman and Reingold 1991).

### qPCR

Bacteria were grown on charcoal agar for 3 days at 37 C before inoculation into Stainer-Scholte (SS) broth (Stainer and Scholte 1970) and grown overnight at 37 C with shaking at 180 rpm; these cultures were used to inoculate fresh media at an OD_600_ = 0.2. Bacterial cells were harvested (1ml for DNA and 10ml for RNA extraction) at OD_600_ = 1.1±0.1 by centrifugation (4000×g for 10 min) and resuspended in 700 μl of Tri-reagent (Invitrogen, ThermoFisher, Loughborough, UK), vortexed vigorously, and frozen at −80°C. DNA was purified using QIAamp kit (Qiagen, Manchester, UK) in accordance with the manufacturer’s instructions. The concentration of DNA was determined using Qubit broad range DNA quantification kit (Fisher Scientific).

qPCR was run on a StepOne Real-time PCR System (Applied Biosystems, ThermoFisher) using TaqMan™ Universal PCR Master Mix (Applied Biosystems), in a total reaction volume of 20 μl with 100pmol of DNA and with primer and probe concentrations as described in Supplement_Tables_7. Triplicate reactions were run for each sample. Reaction conditions were: 10 min at 95°C followed by 40 cycles of 15 sec at 95°C and 1 min at 60°C. Copy number was quantified by using the 2^−ΔΔ*CT*^ method. Three biological repeats were used for determination of copy number in UK54.

To isolate RNA, nucleic acids were precipitated with ethanol, residual DNA was removed by incubation with 4U of Turbo DNase (Ambion, ThermoFisher) for 1 hour at 37 °C, and RNA was purified using the RNeasy kit (Qiagen, Manchester, UK) in accordance with the manufacturer’s instructions. The concentration of RNA was determined using Qubit broad range RNA quantification kit (Fisher Scientific). RNA integrity was determined by agarose gel electrophoresis. Finally, RNA was confirmed as being DNA free by PCR using 50 ng of RNA as template in PCR with recAF and recAR primers. First strand cDNA was synthesised using ProtoScript II (NEB) with 1μg of total RNA as template and 6 μM random primers and incubated for 5 min at 25°C, 1 h at 42°C. The reaction was stopped by incubating at 65°C for 20 min. cDNA was diluted 1/30 in H_2_O for use in qPCR.

RT-qPCR was run on an a StepOne Real-time PCR System using SyberGreen Turbo Master mix (Applied Biosystems), in a total reaction volume of 25 μl with primers at 300 nM. Triplicate reactions were run for each sample. Reactions conditions were: 95°C for 10 min and 40 cycles of 95°C for 15sec and 1 min at 60°C. The housekeeping gene *recA* was used as a stably expressed control gene (Supplement_table_7). The ∆CT and ∆∆CT were calculated by determining the difference between the reference condition and experimental condition. Relative expression was represented as fold change (fold change =2^−∆∆CT^). Significance was determined with one-way ANOVA.

### Electronic mapping

Genomic DNA isolation from *Bordetella pertussis* strains D236, D800, H624, J085, J196, and J321 was performed at the CDC according to a Nabsys solution-based protocol modified from the bacterial DNA protocol for AXG 20 columns and Nucleobond Buffer Set III (Macherey-Nagel, Bethlehem, PA). Purified DNA was sent to Nabsys for nicking, tagging, coating and data collection on an HD-Mapping instrument. Nicking enzyme Nb.BssSI (NEB) was used for strain D236 and the nicking enzyme combination Nt.BspQI/Nb.BbvCI (NEB) was used for strains D800, H624, J085, J196, and J321. Resulting *de novo* assembled HD maps, raw data, and data remapped to PacBio *de novo* assemblies were provided by Nabsys for further analysis and sequence assembly comparisons at the CDC using NPS analysis (v1.2.2424) and CompareAssemblyToReference (v1.10.0.1).

### Nanopore sequencing

*B. pertussis* strain UK54 bacteria were stored at −80°C in PBS/20% glycerol at the University of Bath. Bacteria were grown for 72 hours at 37°C on charcoal agar (Oxoid) plates. Harvested cells were resuspended in 10 ml SS broth to an OD_600_ of 0.1 and grown overnight. At approximately OD_600_ 1.0, cultures were diluted in 50 ml SS broth to an OD_600_ of 0.1 and grown to OD_600_ 1.0. Bacteria were centrifuged at 13 000×g for 5 minutes and processed for gDNA extraction using the protocol available from dx.doi.org/10.17504/protocols.io.mrxc57n. The rapid adaptor (SQK-RAD004) Nanopore library preparation steps were included, adapted for sequencing of very long gDNA molecules.

DNA was sequenced for 48 hours on GridION or MinION sequencers using R9.4 flow cells. Base-calling was performed with Guppy (V2.1.3 or V3.2.1) using the “fast” Flip-flop model. Reads spanning the CNV locus were identified using Blastn alignment with a minimum query length coverage of 90% for the 16kb CNV locus and 10% for the single copy flanking regions (~1kb).

### Identification of structural variants from Nanopore sequence reads

Nanopore reads from UK54 were assembled as previously described (Ring et al. 2018), producing a closed genome of length 4.1Mb, without resolution of any predicted CNVs. To investigate individual reads for putative CNVs or inversions Blastn was used. In order to ease the interpretation of Blastn alignments, the assembly was depleted of homologous regions (e.g., IS481 insertions, rRNA operons) using a 1 kb sliding window, with 200bp step size, and removing all 1kb windows which shared at least 50% homology. The resulting modified assembly was 3.4 Mb (82.9% of full length).

During Nanopore sequencing it is possible for two reads to pass consecutively through a single pore and analysed as a single ‘chimeric’ read-potentially causing false positive CNVs in that read. Because DNA fragments are ligated to adapters during sequencing, such chimeras include an adapter sequence in the middle. Porechop was used to trim adapter sequences from reads (available: https://github.com/rrwick/Porechop). Adapters detected in the middle of reads were trimmed using a lower ‘middle threshold’ identity (75%) rather than default (85%) to ensure a low level of chimeric reads. Each read was aligned to the modified assembly using Blastn and analysed for aligned regions proximal on the read but not on the assembly. Reads of interest were mapped back to the full consensus sequence to analyse the relationships between the DNA ‘junctions’ (the point at which the seemingly disparate regions joined together) and repetitive sequences. Reads which contained a join between two different loci were discarded if the join did not occur in a repeat region (IS, gene duplicate or rRNA). Results were plotted in R using the Circlize package (Gu et al. 2014).

### Association of CNVs with repeat regions

The association of CNV loci with repetitive sequences was tested. Closed genome sequences for isolates containing putative CNVs (excluding previously verified CNVs) were downloaded and the association of CNV boundaries with repeat sequences was determined using R and blast.

### Phylogenetics

To investigate the phylogenetic relationship between strains containing CNVs, a core genome SNP alignment was created using Snippy (available: https://github.com/tseemann/snippy). A phylogenetic tree was constructed using Fasttree (Price et al. 2010) and Itol (Letunic and Bork 2007) was used to display the tree. Ancestral state reconstruction was undertaken in R using the Ace function in the Ape package (Paradis et al. 2004).

### Data Access

All data generated during this study are included in this published article and its supplementary information files. Illumina data for the 28 resolved genome strains is available on the SRA (https://www.ncbi.nlm.nih.gov/sra) with accession numbers SRR9123572, SRR5829828, SRR9006092-3, SRR9123574, SRR9151823, SRR9006149, SRR5829737, SRR5829824, SRR5829749, SRR5829769/SRR5829803, SRR9006067, SRR9118395, SRR5829789/SRR5829798, SRR9118319, SRR9118314, SRR9118293, SRR9118269, SRR9118452, SRR5070923/SRR5514663, SRR5071090/SRR5514664, SRR9131605, SRR9131607, SRR9131663-5.SRR9151824.

Nanopore data is available from NCBI with accession number PRJNA604974. Code needed to reproduce the analysis in this study is contained on Github and is linked to throughout the text.

## Acknowledgements

We thank Josh Quick and Nick Loman, Institute of Microbiology and Infection, School of Biosciences, University of Birmingham for technical assistance with long read sequencing on the Nanopore platform. The data analysis performed here would not have been possible without access to the bioinformatics resource, CLIMB (developed by the MRC, grant number MR/L015080/1). This work was made possible through support from CDC’s Advanced Molecular Detection (AMD) program. The findings and conclusions in this report are those of the authors and do not necessarily represent the official position of the Centers for Disease Control and Prevention.

